# Restoration of BDNF-TrkB signaling rescues deficits in a mouse model of SCA6

**DOI:** 10.1101/2021.02.21.432180

**Authors:** Anna A. Cook, Sriram Jayabal, Jacky Sheng, Eviatar Fields, Tsz Chui Sophia Leung, Sabrina Quilez, Eileen McNicholas, Lois Lau, Alanna J. Watt

**Affiliations:** Biology Department, McGill University, Montreal, QC, Canada; Integrated Neuroscience Program, McGill University, Montreal, QC, Canada; Dept. of Neurobiology, Stanford School of Medicine, Stanford, CA, U.S.A.

## Abstract

Spinocerebellar ataxia type 6 (SCA6) is a neurodegenerative disease resulting in motor coordination deficits and cerebellar pathology. Expression of brain-derived neurotrophic factor (BDNF) is reduced in several neurodegenerative diseases, including in post-mortem tissue from SCA6 patients. Here, we show that cerebellar BDNF levels are reduced at an early disease stage in a mouse model of SCA6 (SCA6^84Q/84Q^). One month of voluntary exercise was sufficient to elevate BDNF expression, as well as rescue both motor coordination and cerebellar Purkinje cell firing rate deficits. A BDNF mimetic, 7,8-dihydroxyflavone (7,8-DHF) likewise improved motor coordination and reversed Purkinje cell firing rate deficits, suggesting that exercise acts via BDNF-TrkB signaling. Prolonged chronic 7,8-DHF administration rescued ataxia when treatment commenced near disease onset, but was ineffective when treatment was started late. These data suggest that 7,8-DHF, which is orally bioavailable and crosses the blood-brain barrier, is a promising therapeutic for SCA6 and argue for the importance of early intervention for SCA6.

## Introduction

Spinocerebellar ataxia type 6 (SCA6) is a rare neurodegenerative disorder characterised by impaired motor coordination and cerebellar pathology. SCA6 is caused by a CAG-repeat expansion mutation in the *CACNA1A* gene^1^. This leads to a progressive loss of motor coordination, which typically onsets in middle age^2^. There is presently no treatment for SCA6^2^, making an understanding of its pathophysiology and identification of novel treatment strategies a high priority. We have used a mouse model of SCA6 to characterise disease pathophysiology. The SCA6^84Q/84Q^ model has a CAG-repeat insertion (84 repeats) in the *CACNA1A* gene^3^. We have previously shown that SCA6^84Q/84Q^ mice develop motor coordination deficits at 7 months^4^, consistent with the mid-life onset of motor dysfunction in SCA6 patients. Interestingly, Purkinje cell loss occurs long after disease onset in SCA6^84Q/84Q^ mice, at 2 years^4^, reminiscent of Purkinje cell loss observed in post-mortem tissue from SCA6 patients^2^. The long latency between the onset of motor deficits and Purkinje cell loss in mice suggests that early motor coordination deficits in SCA6 patients might arise from altered cerebellar function, rather than from cell death, and thus might be reversible if cerebellar function can be restored.

Neurotrophins are molecules that promote neuronal health and survival in both the developing and the adult brain^5^. Interestingly, a reduction in the neurotrophin brain-derived neurotrophic factor (BDNF) has been observed in post-mortem SCA6 cerebellum^6^, which made us wonder whether BDNF signaling deficits contributed to SCA6 pathology. Indeed, a reduction in BDNF expression is observed in many neurodegenerative diseases, including Huntington’s Disease^7^, which like SCA6 is caused by a triplet-repeat expansion mutation, Parkinson Disease (PD)^8^, and Alzheimer Disease (AD)^9^. Additionally, reduced BDNF levels have been reported in several mouse models of ataxia, including SCA1^10,11^, Stargazer^12^, Lurcher^13^, and Purkinje cell degeneration (pcd)^13^. BDNF signals via two receptors: the high-affinity TrkB receptor, that has been implicated as the disrupted signaling pathway in other disease models^14^, as well as the low-affinity p75 receptor.

To explore whether BDNF signaling deficits contribute to SCA6 pathophysiology, we measured expression levels of both BDNF and its high-affinity receptor, tropomyosin receptor kinase B (TrkB) in SCA6^84Q/84Q^ mice. We found that BDNF and its TrkB receptor were both reduced in the cerebellum at the age of onset of motor coordination deficits. Exercise has been shown to elevate BDNF levels in other brains regions in healthy animals^15–18^ and in mouse models of AD^14,19,20^. We found that chronic voluntary exercise partially restored cerebellar BDNF levels, as well as rescuing motor coordination deficits. Interestingly, Purkinje cell firing deficits, which we have previously reported in SCA6^84Q/84Q^ mice^21^, were also reversed, suggesting that Purkinje cell firing deficits can be used as a read-out of cerebellar function in ataxia^22^. To test whether cerebellar BDNF signaling mediated the effects of exercise on ataxia in SCA6, we used a TrkB agonist, 7,8-dihydroxyflavone (7,8-DHF) to mimic BDNF signaling. We found that the BDNF mimetic recapitulated the therapeutic effects of exercise in SCA6^84Q/84Q^ mice, and that this improvement in cerebellar function was associated with the reversal of Purkinje cell firing rate deficits. Furthermore, since the BDNF mimetic we used is orally bioavailable and crosses the blood brain barrier, its use represents a promising novel therapeutic strategy for SCA6 and other forms of ataxia associated with reduced cerebellar BDNF.

## Results

### SCA6^84Q/84Q^ mice display deficits in cerebellar BDNF-TrkB signaling

Deficits in BDNF signaling have been characterised in multiple neurological diseases^7–11,14,19^. Reduced BDNF levels have been observed in tissue from post-mortem human SCA6 cerebellum^6^, representing an advanced stage of disease progression. We wondered whether BDNF is altered at earlier disease stages, and if altered BDNF levels contribute to disease pathology. To address this question, we used a mouse model of SCA6 with a hyper-expanded triplet repeat, SCA6^84Q/84Q^ mice^3^, which allow us to study early disease stages at the time of onset of motor deficits.

We performed immunohistochemistry for BDNF on sections of cerebellar vermis from litter-matched wildtype (WT) and SCA6^84Q/84Q^ mice at 7 months, an age at which we have previously observed the onset of motor deficits^4^. BDNF immunoreactivity was observed in all layers of the cerebellar cortex (**Fig. 1a**). To determine whether BDNF was expressed in neurons or glia, we co-stained for calbindin and GFAP to label Purkinje cells and Bergmann glia respectively. We observed a high degree of colocalization between BDNF and calbindin, but little between BDNF and GFAP, indicating that the BDNF signal in the molecular layer reflects localization predominantly in Purkinje cell dendrites (**Fig. 1a**). We compared BDNF levels in SCA6^84Q/84Q^ and WT mice and found that BDNF immunoreactivity was significantly reduced in cerebellar slices from SCA6^84Q/84Q^ mice in all three cortical layers (**Fig. 1b-d**). BDNF is known to be highly expressed in the hippocampus, and BDNF deficits are observed in the hippocampus in both AD and Down syndrome^14,19,20^, so we wondered whether hippocampal BDNF levels were altered in SCA6. We compared BDNF intensity in hippocampal sections from SCA6^84Q/84Q^ and WT mice and observed no significant differences (**Supplementary Fig. 1**). This suggests that BDNF alterations in SCA6^84Q/84Q^ mice occur predominantly in the cerebellum, the primary locus of degeneration in SCA6^23^.

**Figure 1:**
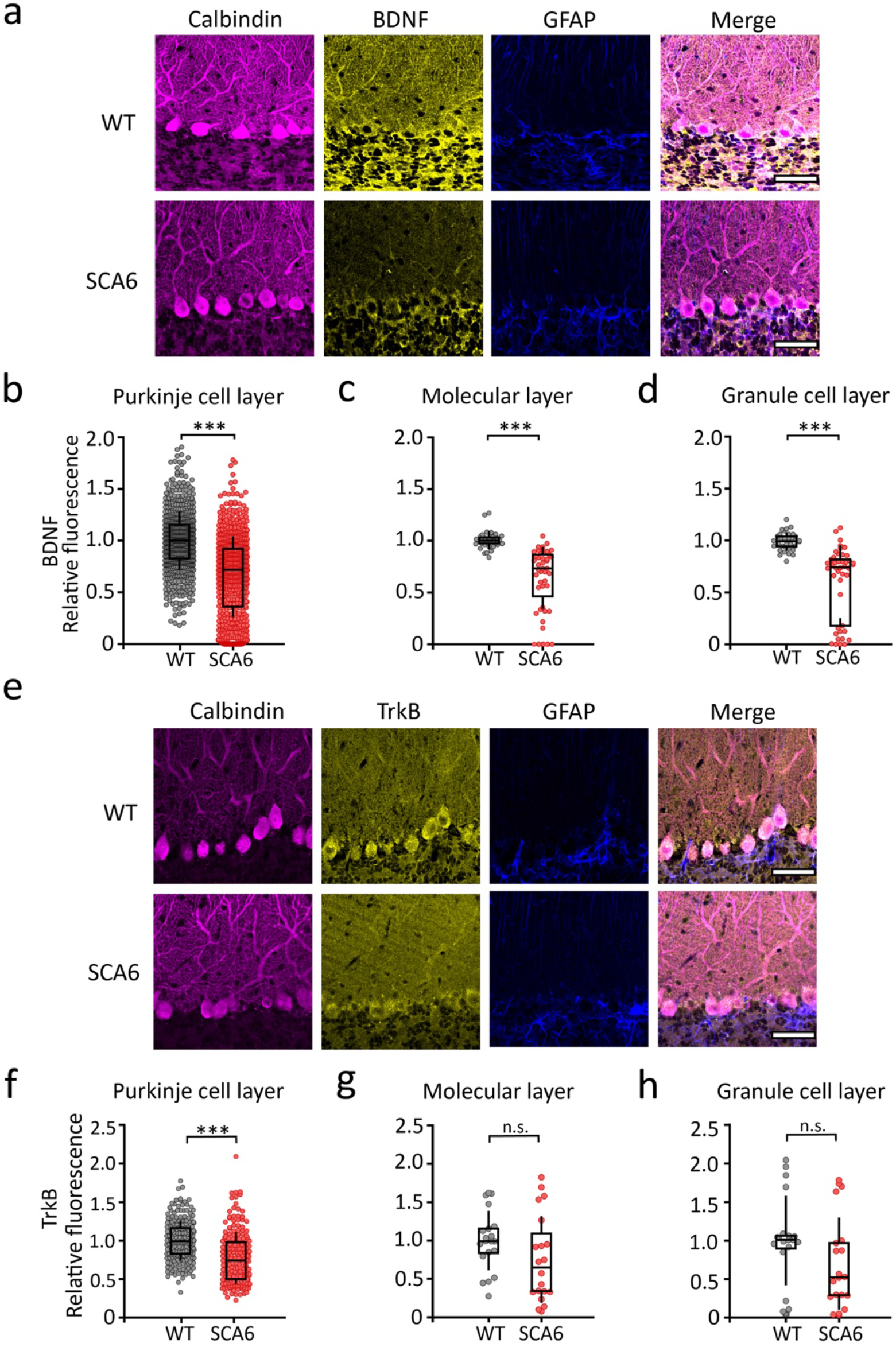
SCA6^84Q/84Q^ mice have reduced cerebellar BDNF and TrkB at disease onset. (**a**) Representative images show BDNF is expressed in all three layers of the WT cerebellar cortex at 7 months but expression is decreased in SCA6^84Q/84Q^ mice. In the upper layers, BDNF colocalizes with Purkinje cells (anti-calbindin) with little labeling in Bergmann glia (anti-GFAP). Scale bar = 50 μm. (**b-d**) Quantification showing BDNF levels are reduced in all the three cortical layers: (**b**) Purkinje cell layer (P<0.0001), (**c**) the molecular layer (P<0.0001), and (**d**) the granule cell layer (P = 0.0002). 4 regions of interest (ROI) were analyzed for each molecular layer and granule cell measurement per animal; WT N = 5 mice, n = 897 Purkinje cell somata; SCA6^84Q/84Q^ N = 6 mice, n = 912 Purkinje cell somata. (**e**) However, TrkB immunoreactivity is reduced only in Purkinje cell somata in SCA6^84Q/84Q^ mice. Scale bar = 50 μm. (**f-h**) Quantification of TrkB expression in the three cortical layers. (**f**) TrkB is reduced in the Purkinje cell layer, but not (**g**) the molecular layer (not significantly different, Students *t* test, P = 0.12), or (**h**) the granule cell layer (P = 0.12). WT N = 4 mice, n = 167 Purkinje cell somata, SCA6^84Q/84Q^ N = 4 mice, n = 125 Purkinje cell somata; Mann Whitney *U* test unless otherwise indicated (when data were normally distributed). ***P < 0.001, n.s. P > 0.05.

We investigated the localization and expression levels of the BDNF receptor TrkB by staining cerebellar vermis sections for TrkB, calbindin and GFAP. Similar to BDNF expression, we observed TrkB immunoreactivity in all cerebellar cortical layers, with the strongest signal arising from Purkinje cell bodies, confirmed by colocalization with calbindin (**Fig. 1e**). TrkB expression in the Purkinje cell and molecular layer colocalized with calbindin but not GFAP (**Fig. 1e**), suggesting that TrkB is expressed in Purkinje cell dendrites and somata. We observed that TrkB staining was reduced in Purkinje cell somas in SCA6^84Q/84Q^ mice compared to WT controls, but was not significantly altered in the molecular layer or granule cell layer at disease onset (**Fig. 1e-h**). Thus, we found that both BDNF and its receptor TrkB are reduced in the cerebellar vermis of SCA6^84Q/84Q^ mice at 7 months, indicating that a deficit in BDNF signaling is observed in the early stages of disease progression in SCA6^84Q/84Q^ mice.

### Exercise elevates BDNF in the SCA6 ^*84Q/84Q*^ mouse cerebellum

Physical exercise can upregulate BDNF expression in several brain regions^15–18^, including in other neurodegenerative disorders in which BDNF levels are reduced^14,19,20^. We wondered whether exercise could increase BDNF levels in the cerebellum of SCA6^84Q/84Q^ mice. To address this, randomly chosen WT and SCA6^84Q/84Q^ mice were provided with running wheels (or locked wheels as a sedentary control) for 1 month beginning at 6 months, before the onset of motor deficits has been reported^4^ (**Fig. 2a**). Both WT and pre-onset SCA6^84Q/84Q^ mice engaged in running behaviour, although we found that SCA6^84Q/84Q^ mice ran shorter distances, and with lower intensity. We found no significant differences in the number of active hours each day between WT and SCA6^84Q/84Q^ mice (**Supplementary Fig. 2**).

**Figure 2:**
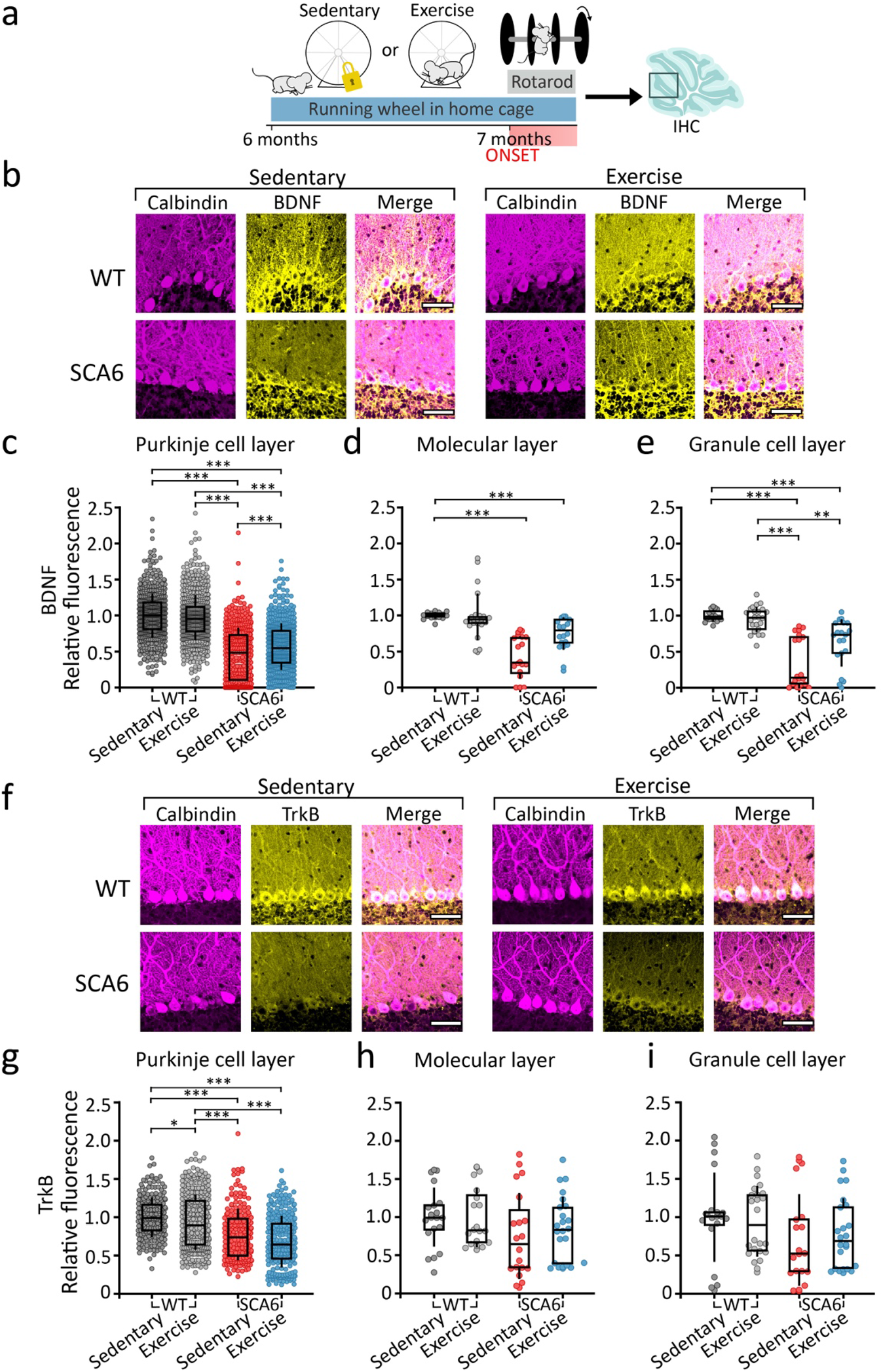
Exercise increases cerebellar BDNF levels in SCA6^84Q/84Q^ mice. (**a**) Schematic of exercise protocol. (**b**) BDNF immunoreactivity in sedentary (left) and exercise (right) WT (top) and SCA6^84Q/84Q^ (SCA6, bottom) mice at 7 months. Scale bars = 50 μm. (**c-e**) Cerebellar BDNF expression in cortical layers: (WT N = 3 mice, WT exercise N = 3 mice, SCA6 N = 3 mice, SCA6 exercise N = 4 mice) (**c**) Purkinje cell layer (All comparisons P <0.0001 except WT Sedentary vs WT Exercise P = 0.45 and SCA6 Sedentary vs SCA6 Exercise, P = 0.0016); (**d**) the molecular layer (All comparisons P <0.005 except WT Sedentary vs WT Exercise, P = 0.41; WT Sedentary vs SCA6 Exercise, P = 0.022; WT Exercise vs SCA6 Sedentary, P = 0.0063; WT Exercise vs SCA6 Exercise, P = 0.27); and (**e**) the granule cell layer (WT Sedentary vs Exercise, P > 0.99; WT Sedentary vs SCA6 Sedentary, P <0.0001; WT Sedentary vs SCA6 Exercise, P = 0.0008; WT Exercise vs SCA6 Sedentary, P = 0.0004; WT Exercise vs SCA6 Exercise, P = 0.0063; SCA6 Sedentary vs SCA6 Exercise, P>0.99). (**f**) Cerebellar TrkB expression in Sedentary (left) and Exercised (right) WT (top) and SCA6^84Q/84Q^ (bottom) mice at 7 months. Scale bars = 100 μm. (**g-i**) Quantification of changes in TrkB expression. (WT N = 4 mice, WT exercise N = 6 mice, SCA6 N = 4 mice, SCA6 exercise N = 6 mice) (**g**) Purkinje cell layer (P < 0.0001 for all comparisons except WT Sedentary vs WT Exercise P = 0.016, SCA6 Sedentary vs SCA6 Exercise P = 0.14); (**h**): molecular layer (P > 0.99 for all comparisons except WT Sedentary vs SCA6 Sedentary, P = 0.25; WT Exercise vs SCA6 Sedentary, P = 0.50), and (**i**) the granule cell layer (P > 0.99 for all comparisons except WT Sedentary vs SCA6 Sedentary, P = 0.31; WT Exercise vs SCA6 Sedentary, P = 0.30). All data is analyzed with one-way ANOVA followed by Dunn’s multiple comparisons test. * P < 0.05, ** P < 0.01, ***P < 0.005, P > 0.05 when comparison is not shown.

To determine whether cerebellar BDNF levels were altered by exercise, we measured BDNF staining intensity in cerebellar vermis of WT and SCA6^84Q/84Q^ mice that were either sedentary or exercised (**Fig. 2b**). We found that exercise elevated BDNF levels in SCA6^84Q/84Q^ mice in Purkinje cell somas (**Fig. 2c**; P = 0.0016), but not in the molecular and granule cell layers (**Fig. 2d,e**). Interestingly, exercise did not alter BDNF levels in WT mice (**Fig. 2c-e**), which is consistent with previous reports^24–26^, and argues that chronic exercise increases cerebellar BDNF expression only when it is pathologically reduced, and that levels are likely saturated in WT mice. Thus, although SCA6^84Q/84Q^ mice run less than WT mice when given free access to a running wheel, they exercise at sufficient levels to alter cerebellar BDNF levels.

Since expression of the BDNF receptor TrkB is also reduced in SCA6^84Q/84Q^ mice (**Fig. 1f**), and because TrkB expression is itself influenced by its ligand BDNF^27^, we wondered whether its expression was also modulated by exercise. We found that TrkB levels were not significantly different in exercised or sedentary SCA6^84Q/84Q^ mice (**Fig. 2g-j**; Purkinje cell layer P = 0.14, molecular layer P > 0.99, granule cell layer P > 0.99), although we saw a modest reduction in TrkB receptor expression in Purkinje cell soma in WT mice (**Fig. 2g**; P = 0.016). This suggests that in SCA6^84Q/84Q^ mice, exercise enhances cerebellar BDNF expression without affecting expression of its high-affinity TrkB receptor.

### Exercise alleviates motor coordination and Purkinje cell firing deficits in SCA6^*84Q/84Q*^ mice

Exercise has been shown to rescue deficits in an SCA1 mouse model^28,29^ and in ataxic Snf2h-null mice^30^, although BDNF has not been identified as the mechanism underlying its therapeutic action in these models. We wondered whether exercise, perhaps via elevation of cerebellar BDNF levels, would rescue motor behavioural deficits in SCA6^84Q/84Q^ mice. To test this, we used an accelerating rotarod to assay motor coordination (**Fig. 3a**). We found that motor coordination in SCA6^84Q/84Q^ mice was significantly improved after 1 month of exercise, compared to sedentary SCA6^84Q/84Q^ mice (**Fig. 3b**), although not to WT levels (**Fig. 3b**). Interestingly, consistent with the lack of change in BDNF expression levels, WT mice showed no significant changes with exercise compared to sedentary controls (**Fig. 3b**). This shows that in addition to restoring cerebellar BDNF levels, exercise reduces ataxia in SCA6^84Q/84Q^ mice.

**Figure 3:**
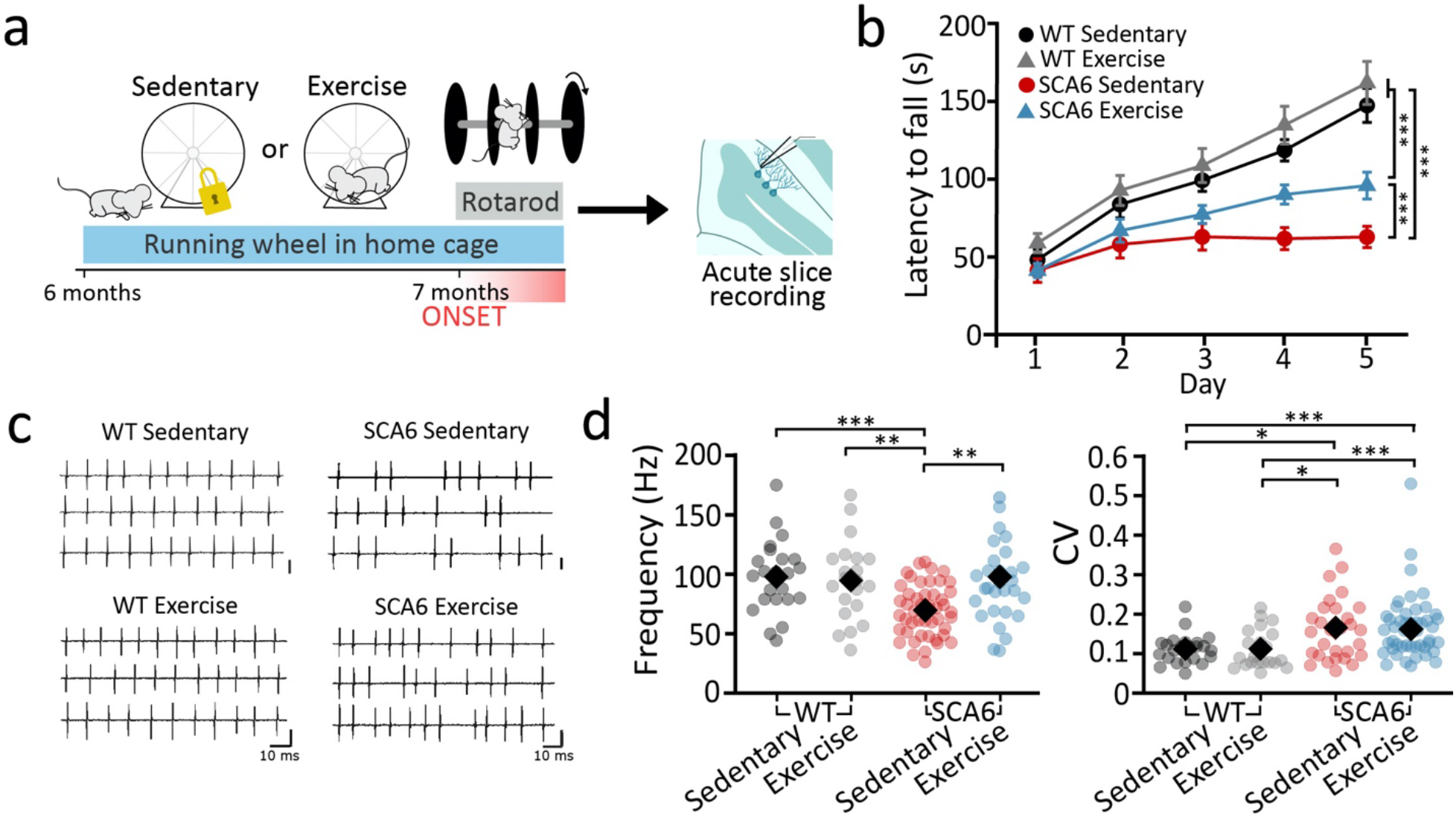
Exercise rescues motor coordination and Purkinje cell firing deficits in SCA6^84Q/84Q^ mice. (**a**) Schematic of exercise protocol. (**b**) Motor coordination is assessed with an accelerating rotarod assay after 1 month of voluntary wheel running by measuring the latency to fall for 5 days of testing (one-way ANOVA followed by Tukey’s multiple comparisons test, P < 0.0001 for all comparisons except WT Sedentary vs Exercise, P = 0.4147 and SCA6 Sedentary vs Exercise P = 0.0002; WT Sedentary, N = 10; WT Exercise, N = 13, SCA6 Sedentary, N = 13; SCA6 Exercise, N = 20). (**c**) Sample juxtacellular Purkinje cell recordings from WT (left) and SCA6^84Q/84Q^ (right) Sedentary (top) or exercise (bottom) mouse cerebellar acute slices. (**d**) Purkinje cell firing frequency (left) is restored in SCA6^84Q/84Q^ Exercise mice (one-way ANOVA followed by Tukey’s multiple comparisons test, WT Sedentary vs WT Exercise, P = 0.98, WT Sedentary vs SCA6 Sedentary, P = 0.001; WT Sedentary vs SCA6 Exercise, P = 0.90; WT Exercise vs SCA6 Sedentary, P = 0.0062, WT Exercise vs SCA6 Exercise P > 0.99; SCA6 Sedentary vs Exercise, P = 0.0055). However, action potential regularity (as measured by the coefficient of variation, CV, of interspike intervals) is not restored by Exercise (WT Sedentary vs Exercise, P = 0.99; WT Sedentary vs SCA6 Sedentary, P = 0.011; WT Sedentary vs SCA6 Exercise, P = 0.0026; WT Exercise vs SCA6 Sedentary, P = 0.014; WT Exercise vs SCA6 exercise, P = 0.0036, SCA6 Sedentary vs Exercise, P = 0.798). n = 29 cells from N = 3 mice WT Sedentary mice; n = 21 cells from N = 3 mice WT Exercise mice; n = 47 cells from N = 3 mice SCA6 Sedentary mice; n = 29 cells from N = 3 mice SCA6 Exercise mice. *P < 0.05, ** P< 0.01, *** P < 0.005, P > 0.05 when not shown.

We next wondered whether other cellular deficits in SCA6^84Q/84Q^ mice were altered by exercise. We have previously shown that spontaneous Purkinje cell firing frequency and regularity are reduced at 7 months in SCA6^84Q/84Q^ mice^21^. Similar deficits in intrinsic Purkinje cell firing have been observed in several ataxia models^10,21,31–33^ (summarised in^22^), suggesting that such firing deficits may be a read-out of cerebellar dysfunction in ataxia. Using juxtacellular recordings of spontaneous Purkinje cell firing in acute sagittal slices, we found that exercise rescued Purkinje cell firing frequency deficits in SCA6^84Q/84Q^ mice to levels indistinguishable from WT mice (**Fig. 3c**). However, firing regularity, quantified by the coefficient of variation (CV) of inter-spike intervals, was unaltered and remained significantly higher in exercise SCA6^84Q/84Q^ mice compared to WT controls (**Fig. 3d**). Exercise did not significantly alter firing properties of Purkinje cells in WT mice (**Fig. 3d**), consistent with the finding that BDNF levels and motor coordination are unaltered by exercise in WT mice. Thus, exercise restores Purkinje cell firing rate and motor coordination deficits, perhaps by restoring cerebellar BDNF levels.

### The BDNF mimetic 7,8-DHF improves motor coordination and Purkinje cell firing rate deficits in SCA6^*84Q/84Q*^ mice

If the behavioural rescue we observed in pre-onset mice with exercise was due to enhanced BDNF expression, we wondered whether targeting BDNF-TrkB signaling alone would be sufficient to rescue motor coordination. To test this, we administered a small molecule TrkB agonist, 7,8-dihydroxyflavone^14,34^ (7,8-DHF) in drinking water (**Fig. 4a, b**).

**Figure 4:**
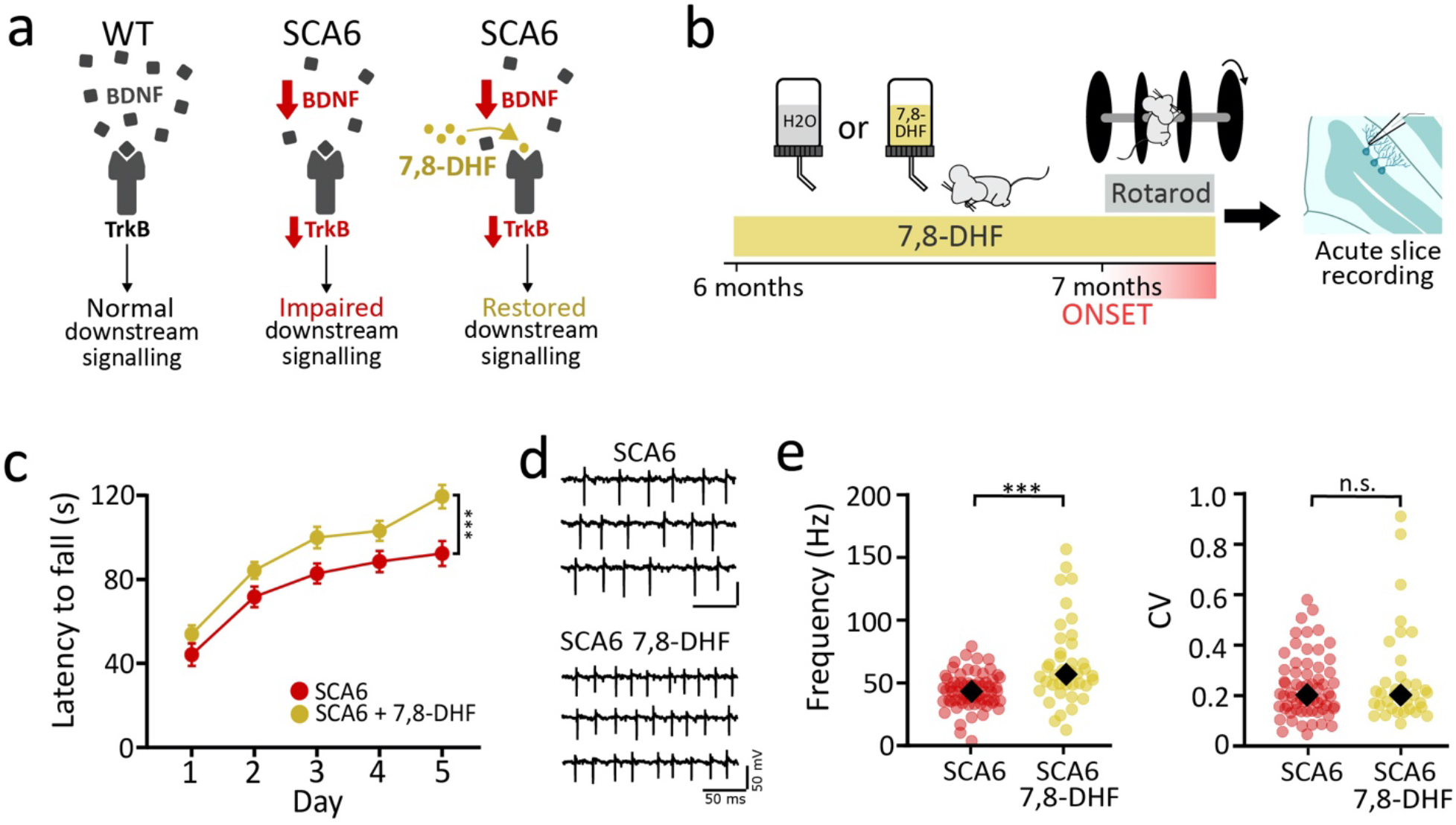
The BDNF mimetic 7,8-DHF restores ataxia and Purkinje cell firing deficits in SCA6^84Q/84Q^ mice at disease onset. (**a**) Proposed model of altered BDNF-TrkB signaling in SCA6^84Q/84Q^ mice and its subsequent restoration with BDNF mimetic 7,8-DHF, a TrkB agonist. (**b**) Schematic of 7,8-DHF administration protocol. (**c**) 7,8-DHF improves motor coordination in SCA6^84Q/84Q^ mice with rotarod assay (SCA6 N = 14 mice, SCA6 + DHF N = 14 mice, significantly different, two-tailed Students *t*-test, P = 0.0014). (**d**) Example traces from juxtacellular recordings of Purkinje cell firing at 7 months after 1 month of 7,8-DHF administration. (**e**) Purkinje cell firing frequency is elevated after 7,8-DHF administration (n = 64 cells from N = 3 mice SCA6 mice; n = 42 cells from N = 3 mice SCA6 + 7,8-DHF, significantly different, Mann Whitney *U* test, P < 0.0001, left). However, Purkinje cell firing regularity is unchanged with 7,8-DHF administration (CV not significantly different, Mann Whitney *U* test, P = 0.86, right). *** P < 0.005, n.s. P > 0.05.

SCA6^84Q/84Q^ mice were given access to drinking water with sucrose and either 7,8-DHF or DMSO vehicle alone as a control at 6 months, prior to onset of motor deficits (**Fig. 4b**), after which we tested motor coordination using the rotarod assay. We found that after 1 month, 7,8-DHF administration alleviates motor coordination deficits in SCA6^84Q/84Q^ mice compared to vehicle controls (**Fig. 4c**). Furthermore, we observed a weak positive correlation between the amount of 7,8-DHF consumed daily and rotarod performance (**Supplementary Fig. 3**), supporting the hypothesis that 7,8-DHF intake influences motor coordination. If 7,8-DHF acts in the cerebellum of SCA6^84Q/84Q^ mice to improve motor coordination, we would predict that it would restore Purkinje cell firing abnormalities^22^. To test this, we carried out juxtacellular recordings of intrinsic Purkinje cell action potentials, and found that similar to the effects of exercise, 7,8-DHF elevated Purkinje cell firing frequency but not firing regularity deficits in SCA6^84Q/84Q^ mice (**Fig. 4d, e**). Thus chronic 7,8-DHF treatment recapitulated the therapeutic effect of the exercise on motor coordination, and also reverses Purkinje cell firing rate deficits that have been previously associated with impaired motor coordination^21,22^. These findings support our hypothesis that exercise and 7,8-DHF act via the same signaling pathway in SCA6^84Q/84Q^ mice.

We hypothesize that WT BDNF levels are near saturation since exercise does not influence motor coordination (**Fig. 3b**), BDNF levels (**Fig. 2d-f**), or Purkinje cell firing properties (**Fig. 3d**) in these mice. We thus expected that 7,8-DHF would not have an effect on motor coordination in WT mice. To test this, we administered 7,8-DHF to WT mice for 1 month, and found it did not significantly affect motor coordination (**Supplementary Fig. 4**). This reinforces our hypothesis that 7,8-DHF affects motor coordination only when cerebellar BDNF levels are reduced.

If exercise restores motor coordination in SCA6^84Q/84Q^ mice through elevation of cerebellar BDNF as we hypothesize, we predict that the combination of exercise and 7,8-DHF treatments should not be additive, whereas if they restore motor coordination through different pathways, combining these treatments would be additive. To differentiate between these possibilities, we compared the effects on ataxia provided by exercise, 7,8-DHF, and the combination of these (**Fig. 5a, b**). We found that exercise alone, exercise + 7,8-DHF and 7,8-DHF alone all significantly rescued motor coordination in SCA6^84Q/84Q^ mice compared to sedentary SCA6^84Q/84Q^ mice, but that there was no significant difference in performance between these three conditions (**Fig. 5c)**. Notably, mice administered Exercise + 7,8-DHF demonstrated no additional improvement in motor coordination in in SCA6^84Q/84Q^ mice compared to either treatment alone (**Fig. 5b**). This supports the hypothesis that exercise improves motor coordination in SCA6^84Q/84Q^ mice by enhancing BDNF-TrkB signaling in the cerebellum. These data suggest that both exercise and 7,8-DHF improve motor coordination via TrkB signaling, and that signaling is near saturation in both conditions.

**Figure 5:**
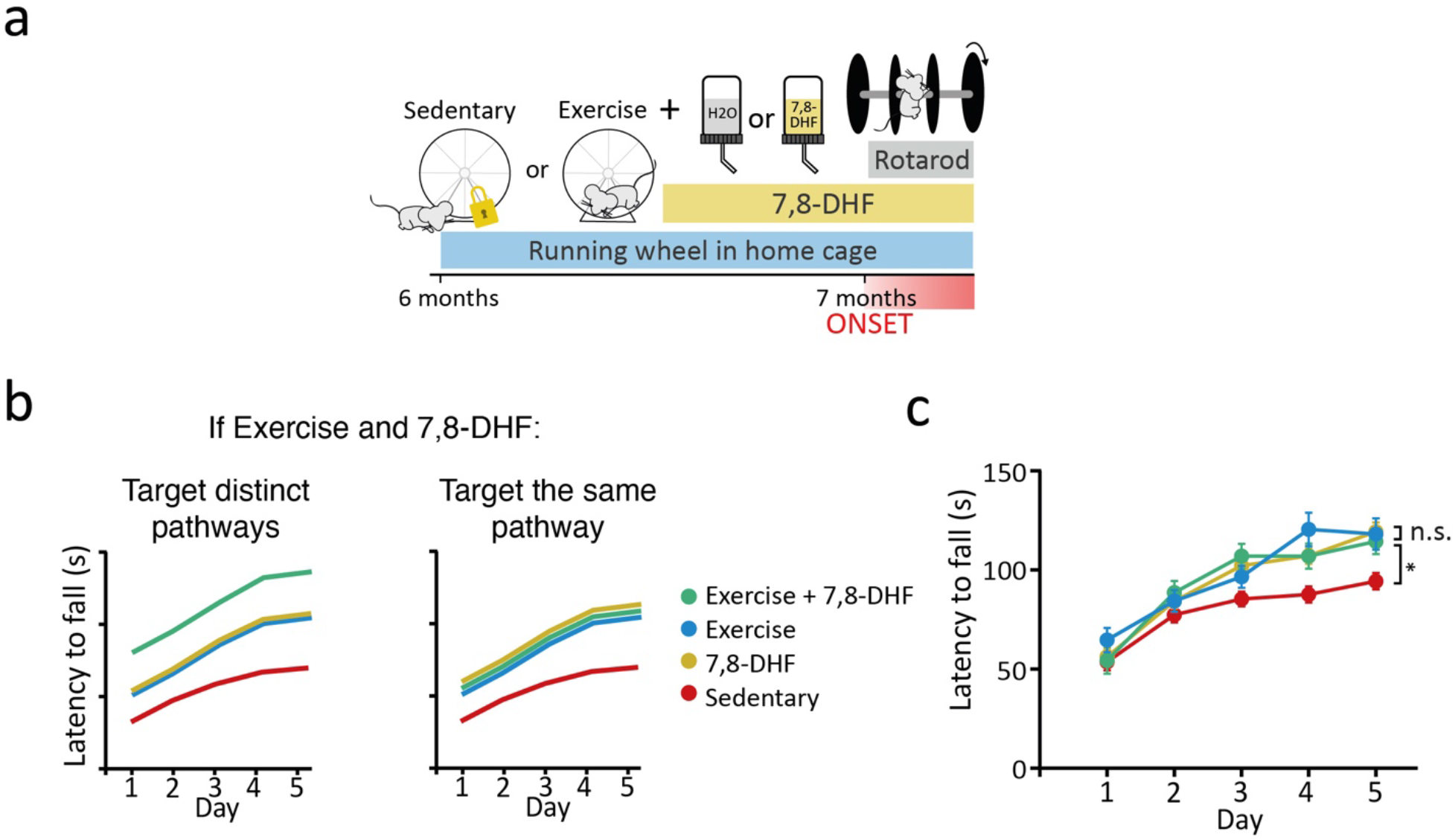
Combining exercise with administration of a TrkB agonist provides no further rescue of ataxia in SCA6^84Q/84Q^ mice. **(a)** Schematic of exercise and 7,8-DHF (DHF) combination therapy on SCA6^84Q/84Q^ mice (SCA6). **(b)** Schematics of possible outcomes if exercise and 7,8-DHF are additive suggesting they act via distinct pathways (left), or if no further improvement is observed in motor coordination when exercise and 7,8-DHF are co-administered, suggesting they act on same signaling pathway (right). **(c)** No additional improvement is observed when exercise and 7,8-DHF administration are combined, and exercise and 7,8-DHF improve motor coordination similarly suggesting they act on the same pathway. (one-way ANOVA followed by Tukey’s multiple comparison test; SCA6 Sedentary vs SCA6 + DHF, P = 0.0009; SCA6 Sedentary vs SCA6 Exercise, P = 0.024; SCA6 Sedentary vs SCA6 Exercise + DHF, P = 0.045; SCA6 + DHF vs SCA6 Exercise + DHF, P = 0.93; SCA6 Exercise vs SCA6 Exercise + DHF, P = 0.98; SCA6 + DHF vs SCA6 Exercise, P > 0.99; SCA6 Sedentary N = 19 mice, SCA6 Exercise N = 12 mice, SCA6 + DHF N = 15 mice, SCA6 Exercise + DHF N = 9 mice). *P < 0.05, n.s. P > 0.05.

### BDNF mimetic 7,8-DHF can restore motor coordination at later disease stages when administration starts early

SCA6 is a progressive disorder which means that motor coordination worsens over time in both patients and mouse models^2,4^. To determine whether targeting TrkB signaling at later ages in SCA6^84Q/84Q^ mice would improve motor coordination, we administered 7,8-DHF at 7 months, after disease onset. SCA6^84Q/84Q^ mice were assayed on rotarod for 5 days and then divided into two equally performing groups, which were given either 7,8-DHF or DMSO vehicle in drinking water (**Fig. 6a**). Rotarod testing continued daily to determine if and when changes in motor coordination occurred. SCA6^84Q/84Q^ mice treated with 7,8-DHF performed significantly better than untreated controls after 14 days, and performance tended to increase for the next 2 weeks in treated mice (**Fig. 6b**). The relative slow time-course of 7,8-DHF action is consistent with our finding that 4-hour 7,8-DHF treatment on acute cerebellar slices was insufficient to alter Purkinje cell firing properties (**Supplementary Fig. 6**), supporting our observation of a slow time course to ameliorate motor coordination (**Fig. 6b**). Our findings that restoring TrkB signaling with 7,8-DHF at the onset of ataxia still rescued motor coordination in SCA6^84Q/84Q^ mice are particularly promising since mice did not exercise at high levels at this age, and exercise was unable to rescue deficits in SCA6^84Q/84Q^ mice when started after onset, even when supplemented with bouts of forced exercise on a treadmill (**Supplementary Fig. 5**). The difference we observe in the efficacy of exercise and 7,8-DHF treatment post-onset may arise from motor impairment due to disease progression that prevents SCA6^84Q/84Q^ mice from exercising at sufficiently high levels.

**Figure 6:**
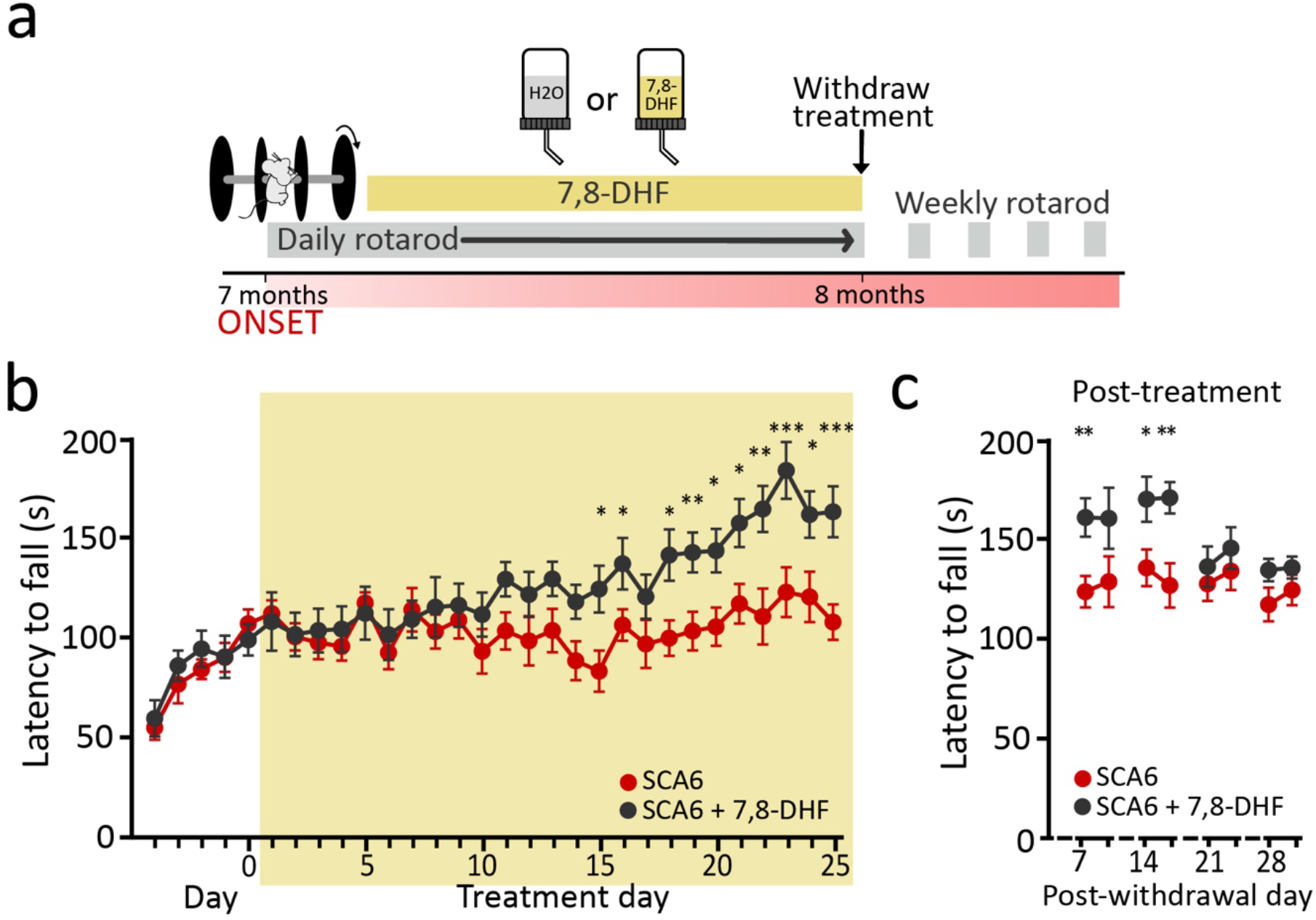
Chronic administration of 7,8-DHF leads to a slow timecourse of improvement of ataxia in SCA6^84Q/84Q^ mice. (**a**) Schematic of 7,8-DHF administration protocol. (**b**) SCA6^84Q/84Q^ mice (SCA6) were administered 7,8-DHF for 25 days after 5 days of baseline. Improvement in motor coordination was observed after ~ 2 weeks (SCA6 N = 4 mice, SCA6 + 7,8-DHF N = 4 mice, all days compared with Mann Whitney *U* tests or Student’s *t* tests, days marked with asterisks were significantly different; treatment day 14, P= 0.030; day 15, P = 0.013; day 18, P = 0.011; day 19, P = 0.0081; day 20, P = 0.013; day 21, P = 0.014; day 22, P = 0.0061; day 23, P = 0.0031; day 24, P = 0.023; day 25, P = 0.0020, all other P values were >0.05). (**c**) Motor performance remained elevated for ~2 weeks after 7,8-DHF withdrawal (SCA6 N = 3, SCA6 DHF (withdrawn) N = 3, all days compared with Mann Whitney *U* tests or Student’s *t* tests, days marked with asterisks were significantly different: day 7, P= 0.0092; day 14, P = 0.011; day 15, P = 0.0062, all other p values were >0.05). * P < 0.05, ** P < 0.01, P > 0.05 when not shown.

Given that BDNF expression is regulated by BDNF-TrkB signaling^27^ we wondered if the effects of 7,8-DHF might persist in SCA6^84Q/84Q^ mice. To test this, we withdrew 7,8-DHF after 1 month while testing motor coordination weekly. We found that 7,8-DHF-treated mice maintained elevated motor coordination for 2 weeks after 7,8-DHF was removed, but that motor coordination returned to control levels by 3 weeks (**Fig. 6c**). Thus, 7,8-DHF treatment had an effect over a slow time course on motor coordination both for drug administration and withdrawal.

SCA6 patients often live with the progressively worsening disease for decades. We wondered whether 7,8-DHF could provide long-lasting benefits in SCA6^84Q/84Q^ mice. To address this, we continued 7,8-DHF treatment for 4 months in a subset of SCA6^84Q/84Q^ mice and assayed motor coordination monthly (**Fig. 7a**). We found that SCA6^84Q/84Q^ mice chronically administered 7,8-DHF showed improved performance at 7, 9, and 10 months (**Fig. 7b**). This suggests that the continued administration of 7,8-DHF may have a long-lasting therapeutic benefit in SCA6 patients. To determine whether chronic 7,8-DHF administration had similar cellular effects in older mice that it did in younger mice, we recorded Purkinje cell firing rates in acute slices made after 4 months of 7,8-DHF and found that they were not significantly different than controls (**Fig. 7c**), unlike what we observed in younger mice where 7,8-DHF restored Purkinje cell firing rate deficits (**Fig. 4c**).

**Figure 7:**
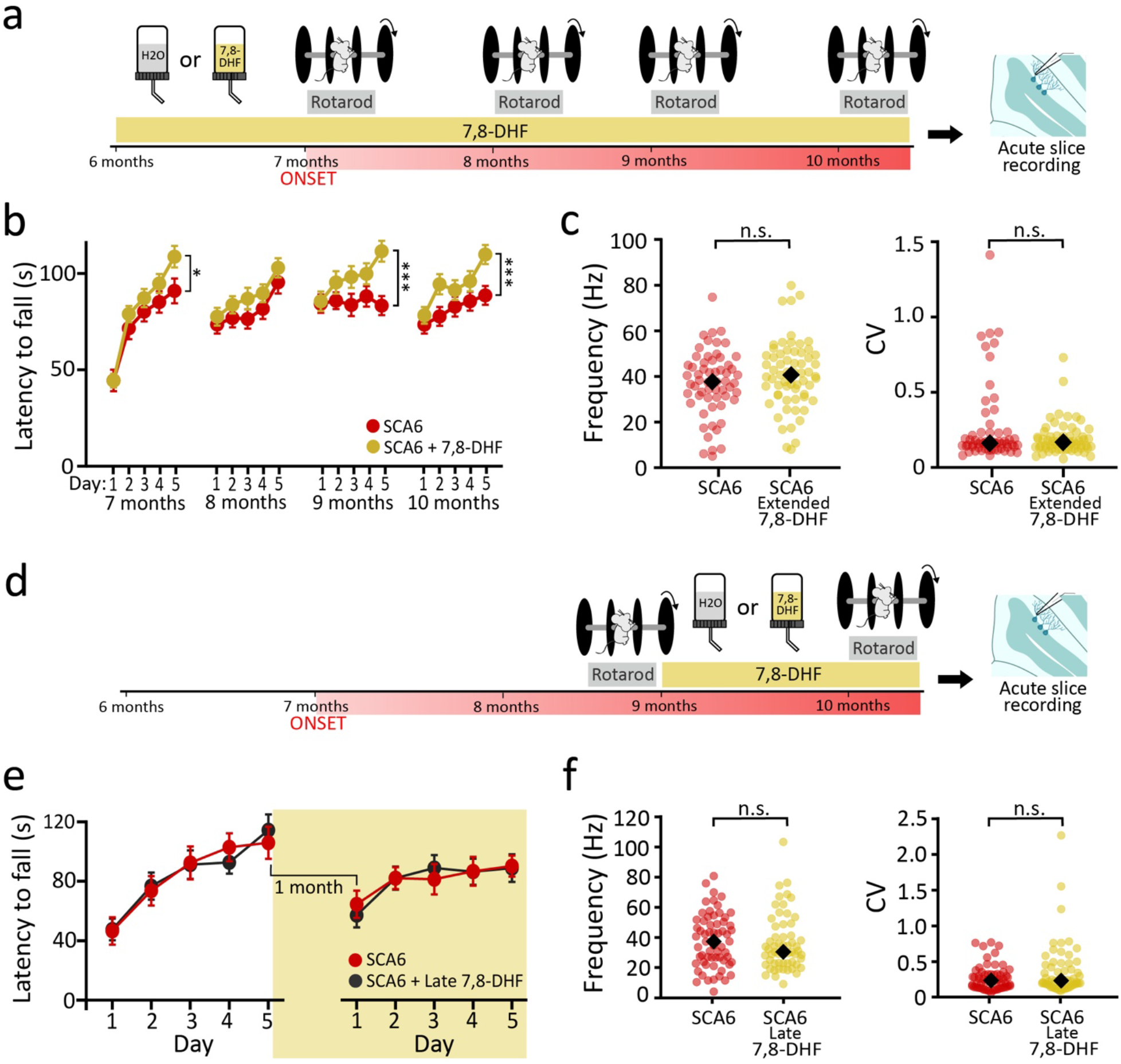
Early, but not late, 7,8-DHF administration rescues deficits as disease progresses in SCA6^84Q/84Q^ mice. (**a**) Schematic of long-term chronic 7,8-DHF (DHF) administration in SCA6^84Q/84Q^ mice started near disease onset. (**b**) Mice treated chronically with 7,8-DHF performed better at 7, 9, and 10 months old (SCA6 N = 11 mice, SCA6 DHF N = 9 mice, all months were compared using T-test, 7 month p = 0.041, 8 month p = 0.33, 9 month p<0.0001, 10 month p = 0.00014) (**c**) Purkinje cell firing rate (left) was no longer restored after 4 months of drug administration (SCA6, n = 58 cells from N = 2 mice; SCA6 DHF n = 62 cells from N = 3 mice; not significantly different, two-tailed Student’s t test, P = 0.22), nor was regularly affected (not significantly different, Mann-Whitney *U* test, P = 0.63). (**d**) Schematic of 7,8-DHF administration started in late disease progression. (**e**) Late-administered 7,8-DHF does not restore firing deficits at 10 months when treatment starts after disease onset at 9 months. (not significantly different before or after DHF administration, two-tailed Student’s *t* tests; before P = 0.91, after P = 0.58; SCA6 N = 5 mice; SCA6 late DHF N = 4 mice). (**f**) Purkinje cell firing rate and regularity (CV) are not significantly different after late administration of 7,8-DHF (left: frequency not significantly different, Mann Whitney *U* test, P = 0.46; right, CV not significantly different, P = 0.17; SCA6, n = 62 cells from N = 3 mice, SCA6 late DHF, n= 59 cells from N = 3 mice). *P < 0.05, *** P < 0.001, n.s. P> 0.05.

Our observation that treatment with the BDNF mimetic 7,8-DHF provides long-term therapeutic benefits in the SCA6^84Q/84Q^ mouse model (**Fig. 7b**) is promising for SCA6 patients. However, SCA6 is often diagnosed after disease onset, meaning that early treatment may not always be practical. We thus wondered whether 7,8-DHF would ameliorate ataxia in SCA6^84Q/84Q^ mice when drug administration was initiated at a later stage of disease progression. To determine whether this was likely to succeed, we first determined that TrkB receptors were still expressed in 9 month SCA6^84Q/84Q^ cerebellum (**Supplementary Fig. 7**), suggesting that the receptor target for 7,8-DHF is expressed in the SCA6 brain. We then administered 7,8-DHF for 1 months to SCA6^84Q/84Q^ mice at 9 months (**Fig. 7d**). We observed no significant improvement in motor coordination in late-treated 7,8-DHF mice (**Fig. 7e**), and similarly, observed no changes in Purkinje cell firing properties (**Fig. 7f**). These data suggest that the lack of a therapeutic effect of late-administered 7,8-DHF is unlikely to arise due to TrkB receptors, but rather likely arises from changes in downstream signaling mechanisms. Taken together, our results demonstrate that 7,8-DHF is a promising therapeutic with the potential for long-lasting benefits for SCA6-associated ataxia when administration is started around disease onset, but not at later stages.

## Discussion

We identified BDNF-TrkB signaling disruptions as a contributor to pathophysiology at early disease stages in SCA6^84Q/84Q^ mice and successfully targeted this pathway to improve disease phenotype using two approaches. Our work represents a promising novel therapeutic strategy for SCA6 and sheds light on the mechanisms underlying the therapeutic properties of exercise for SCA6. Reduced cerebellar BDNF has been reported in post mortem brain tissue from SCA6 patients^6^ and we identified similar changes in the SCA6^84Q/84Q^ mouse model, alongside reduced levels of the BDNF receptor TrkB. We went on to show that cerebellar BDNF could be rescued by a chronic exercise program, which also rescued deficits in motor coordination and Purkinje cell firing. We then used 7,8-DHF, a BDNF mimetic that acts as a TrkB agonist, to rescue deficits in the SCA6^84Q/84Q^ mouse. Combining these two therapies provided no additional rescue of motor coordination deficits, suggesting that exercise exerts its beneficial effects on motor coordination by restoring TrkB signaling in the cerebellum. 7,8-DHF treatment improved motor coordination for months if treatment was initiated around the onset of ataxia, but was insufficient if treatment was started later in the disease progression.

Many studies using animal models of human diseases focus on early, peri-onset disease stages (e.g.^21^). Yet for progressive, neurodegenerative diseases like SCA6, patients often live with symptoms for years or decades, and diagnosis is sometimes made long after disease onset. For this reason, we chose to focus on multiple stages of disease progression: pre-onset, early progression, and late progression. We found that in our mouse model, both exercise and TrkB agonist treatment mitigated the development of motor coordination and Purkinje cell firing deficits when administered pre-onset. Exercise did not contribute further improvement of motor coordination by 7,8-DHF suggesting that exercise and 7,8-DHF act via the same pathway. Later, as disease progressed, exercise ceased to rescue motor function, likely because the mice did not run far enough. Nonetheless, treatment with 7,8-DHF was still able to reverse deficits, demonstrating that restoring BDNF was still possible at early stages of disease progression. This argues that different windows of opportunity may exist for different interventions, although exercise in humans after disease onset may be more successful than it was in our mouse model. Importantly, however, at a more advanced stage later in disease progression, 7,8-DHF treatment was only able to restore motor when administration began peri-onset and continued for 4 months, but not when administration began later. With increased prevalence of predictive genetic testing in neurodegenerative diseases like SCA6^35^, an awareness of the importance of early treatment may be of benefit to patients.

Exercise is an intervention that is accessible to many patients and has been successfully used to improve outcomes for ataxia; for example, exercise leads to some alleviation of motor symptoms and prevention of cerebellar decline in patients with various types of ataxia, including SCA6^36–39^, although until now there was no evidence that this therapeutic action was mediated by BDNF. Exercise has also proven therapeutic in other animal models of ataxia^28–30^, although the mechanism of rescue described in those models appears at least partially different from the mechanism we describe here, perhaps because of differences in the duration and intensity of exercise animals undergo. It is also likely that the difference in the molecular mechanism of therapeutic exercise in different ataxias arises due to differences in the molecular pathophysiology of different disorders. While it is unclear what the equivalent of 30 days of voluntary wheel running by a mouse would be for an SCA6 patient, we know that there are exercise protocols, such as high-intensity interval training, that target BDNF-TrkB signaling in people^40,41^. Thus, despite our findings that exercise ameliorates ataxia may be generalizable, it will be nonetheless important to determine how exercise acts in each form of ataxia, to determine whether a strategy targeting BDNF signaling will be beneficial.

While the upregulation of BDNF with exercise has been well characterised in the hippocampus^42^, where even short bouts of exercise lead to upregulation of BDNF, the picture in the cerebellum has been less clear. Some studies in rodents have reported no change in BDNF expression in the cerebellum with exercise^24–26^, while others report that exercise increases levels of cerebellar BDNF^15,43^. These discrepancies could arise due to differences in the duration and type of exercise in each protocol. We found that although 4 weeks of exercise did not alter cerebellar BDNF protein levels in WT mice, it did increase BDNF protein levels in the SCA6^84Q/84Q^ cerebellum. This differential effect on WT and disease model mice is not unprecedented, and other studies similarly report that exercise acts differently in different models and circumstances^44,45^. We surmise that the chronic exercise protocol that we used increased BDNF levels only when they were already pathologically reduced in the SCA6^84Q/84Q^ mice.

Reduced BDNF expression is a hallmark of many neurological diseases. BDNF is known to be upregulated by neuronal activity, and so the reduction in SCA6 and other ataxias could be a secondary effect caused by the decreased firing frequency of Purkinje cells that has been reported for SCA6^21^ and many other ataxic mouse models^22^. However, other mechanisms leading to reduced BDNF in disease cannot be excluded, and indeed, BDNF disruption may arise from a variety of different insults in different diseases.

The restoration of BDNF-TrkB signaling we describe here for SCA6 is a promising potential therapeutic approach for multiple ataxias. Our results are consistent with previous findings that viral reintroduction of the BDNF gene rescued ataxia in Stargazer mice^12^ and acute cerebellar delivery of BDNF to SCA1 *ATXN1(82Q)* mouse model had a similar therapeutic effect^11,46^. However, overexpression of BDNF has been associated with negative outcomes in some cases^47–49^, likely due to the complex signaling pathways that are known to be activated by BDNF. Thus, instead of virally reintroducing BDNF in the SCA6^84Q/84Q^ mouse, we chose to use a different approach administering a BDNF mimetic that acts as a TrkB agonist, as this signals exclusively through the neuroprotective TrkB pathway and may be more translatable to the clinic^5^.

In summary, we have found that both BDNF and its high-affinity TrkB receptor are downregulated at the age of onset in a mouse model of SCA6. We found that exercise elevated BDNF in the cerebellum of SCA6 mice, and that this was associated with an improvement of motor coordination deficits and cellular firing deficits in cerebellar Purkinje cells. We then used a BDNF mimetic to activate TrkB receptors directly and found this restored motor coordination and Purkinje cell firing deficits in a manner similar to exercise. When administered together, exercise and the BDNF mimetic were not additive, suggesting that they were acting on the same pathway. Chronic administration of the BDNF mimetic that was initiated around the age of disease onset led to a sustained improvement in motor coordination, but this was not true when administration started later during disease progression. Taken together, our data suggest that exercise modulates ataxia in SCA6 through the restoration of BDNF-TrkB signaling in the cerebellum and that a BDNF mimetic can mimic this therapeutically. Our work therefore highlights a promising therapeutic approach to both SCA6 and other neurodegenerative diseases.

## Methods

### Animals

We used a knock-in mouse model of SCA6 with a humanised 84 CAG repeat expansion mutation at the *CACNA1A* locus (SCA6^84Q^). Mice that were heterozygous at this locus were obtained from Jackson laboratories (Bar Harbor, Maine; strain: B6.129S7-Cacna1atm3Hzo/J; stock number: 008683) and were bred to obtain SCA6^84Q/84Q^ and litter-matched WT control mice. Genotyping was performed at weaning using primer sequences published by Jackson laboratories. Male and female mice were used in all experiments. All animal procedures were carried out with the approval of the McGill Animal Care Committee in accordance with the Canadian Council on Animal Care guidelines.

### Immunohistochemistry

Tissue fixation was carried out as previously described^31^. Mice were deeply anesthetised with intraperitoneal injection of 2,2,2-tribromoethanol (Avertin) and an intracardiac perfusion was performed with a flush of ice-cold phosphate-buffered saline (PBS, 0.1M, pH 7.4) with 5.6 μg/mL heparin salt, followed by 40 mL of 4% paraformaldehyde (PFA) in phosphate buffer (PB, pH 7.4). Extracted tissue was stored in 4% PFA at 4°C for a further 24 hours of fixation before being transferred to PBS with 0.5% sodium azide for storage at 4°C. The cerebellar vermis was dissected and sliced into 100 μm sagittal slices with a Vibratome 3000 sectioning system (Concord, ON, Canada). Immunohistochemistry on free floating slices was carried out with antibodies for proteins of interest along with anti-calbindin or anti-GFAP as markers for Purkinje cells and Bergmann glia, respectively. Antibodies are listed in **Supplementary Table 1**. The use of anti-BDNF included an additional heat-induced epitope retrieval step comprising 10 minutes of heating to 95°C in PBS, after which slices were allowed to cool before proceeding with the primary antibody staining. After staining, slices were immediately mounted in ProLong Gold Antifade mounting medium (ThermoFisher Scientific, Waltham, USA) and stored in the dark at 4°C.

### Image acquisition and analysis

Imaging was performed using an LSM800 confocal microscope (Zeiss), using Zeiss Zen software for image acquisition. The raw fluorescence intensity was recorded in arbitrary units and was normalised to the mean WT value within a batch. Due to limitations of litter size, immunocytochemistry was typically carried out in several batches. To minimise batch-specific effects on the data, all data is shown normalised to the mean WT value of that batch (defining WT intensity = 1). Image analysis was performed in FIJI (ImageJ; US National Institutes of Health)^50,51^. Raw images of lobule 3 of cerebellar vermis were used and manual tracing around Purkinje cell bodies in the calbindin channel allowed the delineation of ROIs around the Purkinje cell bodies. Missing or out of the plane of focus cells were skipped. These ROIs were then exported onto the BDNF/TrkB channel and fluorescence intensity was measured (integrated density, arbitrary units). For measurements of fluorescence intensity in the molecular layer and granule cell layer, we used 4 ROIs of consistent size (5193 μm^2^) to determine the total intensity at randomly chosen locations in the molecular and granule cell layers (**Supplementary Fig. 8**).

### Voluntary running

Mice were individually housed in low profile rat cages with running wheels (Columbus Instruments, OH, USA). Sedentary control mice received wheels that were locked in position to control for environmental enrichment provided by the presence of the wheel and the larger cages. The number of wheel rotations was recorded hourly using Multi-Device software (Columbus Instruments, OH, USA).

### Treadmill exercise

Mice were moved to the behavior room for acclimatization one hour prior to the start of exercise. Mice were placed individually onto an enclosed single lane treadmill (Treadmill Simplex II Columbus Instruments). The treadmill speed was set to 6 m/min for the first 5 minutes, then increased to 10 m/min for 5 minutes, then increased in 1 m/min increments for each subsequent 5 minute period of the 20-minute exercise period. The treadmill was equipped with a shock plate, which distributed 2 mA shocks twice per minute when mice stepped off the treadmill. If a mouse remained on the shock plate for more than 30 seconds, the treadmill was stopped for an additional 30 seconds allowing the mouse to return to the treadmill. Mice in the control condition were placed on the stationary treadmill and allowed to explore it for 20 minutes with the shock platform on, encouraging the mice to remain on the treadmill. This exercise protocol was repeated daily for 6 weeks.

### Accelerating rotarod

We used an accelerating protocol on a Rotarod (Stoelting, IITC) as previously described ^4,21,31^. We have previously shown that this assay is able to detect differences in motor coordination in WT and SCA6^84Q/84Q^ mice in both control and treatment conditions^4,21^. Mice were moved to the testing room and left to acclimatise for 1 hour prior to testing. Mice underwent 4 trials per day with a 10-15 minute rest period between trials. Testing was carried out at the same time each day for 5 consecutive days or in some cases for up to 30 days. The protocol started with a speed of 4 RPM and then accelerated to 40 RPM over 5 minutes after which the assay continued at 40 RPM for up to 5 further minutes. Latency to fall off the rotarod was recorded in each trial.

### Acute Slice preparation

Slice preparation was carried out as previously described^21^. Mice were deeply anesthetised with intraperitoneal injection of 2,2,2-tribromoethanol (Avertin) and an intracardiac perfusion was performed with ice cold partial sucrose replacement slicing solution (111 mM Sucrose, 50 mM NaCl, 2.5 mM KCl, 0.65 mM CaCl2, 10 mM MgCl2, 1.25 mM NaH2PO4, 25 mM NaHCO3 and 25 mM glucose, bubbled with 95% O2 and 5% CO2 to maintain pH at 7.3; osmolality ~320 mOsm). The cerebellar vermis was dissected and sliced into 200 μm sagittal slices with a VT 1200S vibratome (Leica Microsystems, Wetzlar, Germany). Slices were incubated in artificial cerebrospinal fluid (ACSF: 125mM NaCl, 2.5mM KCl, 2mM CaCl2, 1mM MgCl2, 1.25mM NaH2PO4, 26mM NaHCO3 and 20mM glucose, bubbled with 95% O2 and 5% CO2 to maintain pH at 7.3; osmolality ~320 mOsm) bubbled with carbogen at 37°C for 45 minutes and then transferred to room temperature for the duration of experiments. All reagents were obtained from Sigma (Oakville, ON, Canada) apart from MgCl_2_ and CaCl_2_ obtained from Fisher Scientific (Toronto, ON, Canada).

### Electrophysiology

Recordings of spontaneous Purkinje cell firing in lobule 3 of cerebellar vermis were carried out as previously described^21,31^. Slices were kept in a bath of ACSF at approximately 33°C during recording. Purkinje cells were visually identified and juxtacellular recordings were made at the soma with borosilicate pipettes pulled with a P-1000 puller with tip sizes of 1-2μm (Sutter Instruments, Novato, CA, USA). Data acquisition and analysis was performed in IGOR Pro version 6.0 and 7.0 (Wavemetrics, Portland, OR, USA) using custom routines.

### 7,8-DHF administration

7,8-dihydroxyflavone (7,8-DHF) is orally bioavailable and previous studies have demonstrated that oral administration via drinking water is both well tolerated and able to deliver therapeutically relevant dosages of the drug in mice^14^. Single-housed mice were given free access to solutions containing either 157.3 μM 7,8-DHF (TCI America) dissolved in 0.04% DMSO or control solutions with 0.04% DMSO only, dissolved in autoclaved water. 10% sucrose was added to both solutions to make them palatable, as previously described^21^. Solutions were made fresh twice weekly and refreshed daily, with solutions stored at 4°C in between refills. Consumption was monitored daily to determine the administered dose of 7,8-DHF. Mice consumed on average 21 mL of the sucrose-sweetened drug solution daily, giving a daily dose of 0.84 mg 7,8-DHF. The control group with 0.04% DMSO and 10% sucrose but no 7,8-DHF consumed on average 19 mL of the solution each day. The drug treatment period ranged from 25 days to 4 months and mice underwent behavioural testing on the rotarod assay during this time. Mice were monitored daily when solutions were administered and no adverse health effects were observed. Mice were weighed at least every second day during behavioural testing, meaning that some mice receiving 7,8-DHF were weighed throughout their drug treatment and did not gain or lose weight during the treatment (**Supplementary Fig. 3**). Reagents were obtained from Sigma (Oakville, ON, Canada) unless otherwise specified.

### Statistics

Comparisons were made using either one-way ANOVA followed by post hoc tests using JMP software (SAS, Cary, NC), or two-tailed Student’s *t* tests or Mann-Whitney *U* tests using Igor Pro software, for parametric or non-parametric testing respectively. Data are reported as Mean ± S.E.M. when normally distributed, or as median when the data are not normally distributed.

## Author Contributions

A.A.C designed and ran experiments and analyzed data for all Figures except Fig. 3 and wrote the manuscript, S.J. designed and ran experiments and analyzed data for Fig. 3 and helped write the manuscript, J.S. designed and ran experiments and analyzed data for Fig. 1 and 2, E.F. ran experiments for Fig. 7 and analyzed data for Fig. 2, T.C.S.L. analyzed data for Supplementary Fig. 7, S.Q. ran experiments and analyzed data for Fig. 3, E. M. ran experiments and analyzed data for Supplementary Fig. 4, L.L. analyzed data for Supplementary Fig. 1, 2 and 3, A.J.W. conceived of the project, designed experiments and analyzed data, supervised the project, and wrote the manuscript.

## Acknowledgements

We thank Anne McKinney, Aparna Suvrathan and Jesper Sjöström for helpful advice and thoughtful feedback on the project. Custom Igor software routines for electrophysiology data acquisition and analysis were written by Jesper Sjöström. We thank Karim Nader and Arkady Khoutorsky for the use of equipment. We thank all current and former members of the Watt lab for their input, particularly Kristen Vieira-Lomasney, Mohini Bhade and Genavieve Maloney who provided technical support, and Brenda Toscano-Márquez, Amy Smith-Dijak, Daneck Lang-Ouellette, Kim Gruver, and Louisa Shen for their helpful feedback. Imaging was performed in the McGill University Advanced BioImaging Facility (ABIF) and we thank ABIF staff members for their technical support. We are grateful for the excellent animal care and training we received from the McGill Animal Resources Centre (CMARC), particularly Tanya Koch and Juan Canale who greatly facilitated the research.

## Supplementary Information

Supplementary Figs. 1-8

Supplementary Table 1

