## Supplementary Figures and Table for "Restoration of BDNF-TrkB signaling rescues deficits in a mouse model of SCA6"

### Exercise acts via BDNF-TrkB signalling to rescue deficits in a mouse model of SCA6

**Supplementary Fig. 1-8**
**Supplementary Table 1**

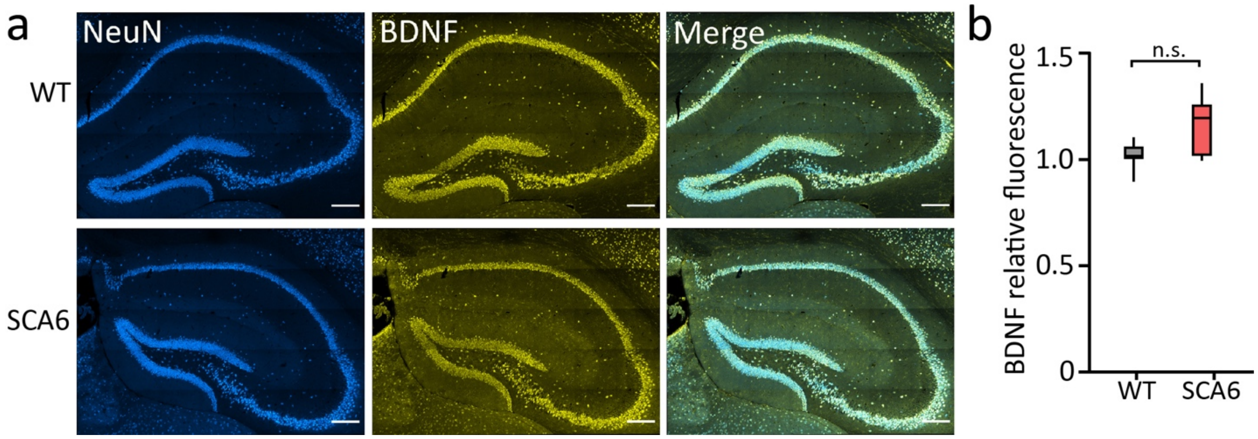


**Supplementary Fig. 1. SCA6$^{84Q/84Q}$ mice do not show changes in hippocampal BDNF levels.**

(**a**) Representative hippocampal slices showing cell bodies (anti-NeuN) and BDNF expression from WT (top) and SCA6 (bottom) mice. Scale bars = 100 µm. (**b**) BDNF intensity in hippocampus is not significantly altered in SCA6 mice (WT N = 3 mice, SCA6 N = 3 mice, not significantly different, Mann Whitney *U* test, P = 0.41). n.s. P > 0.05.

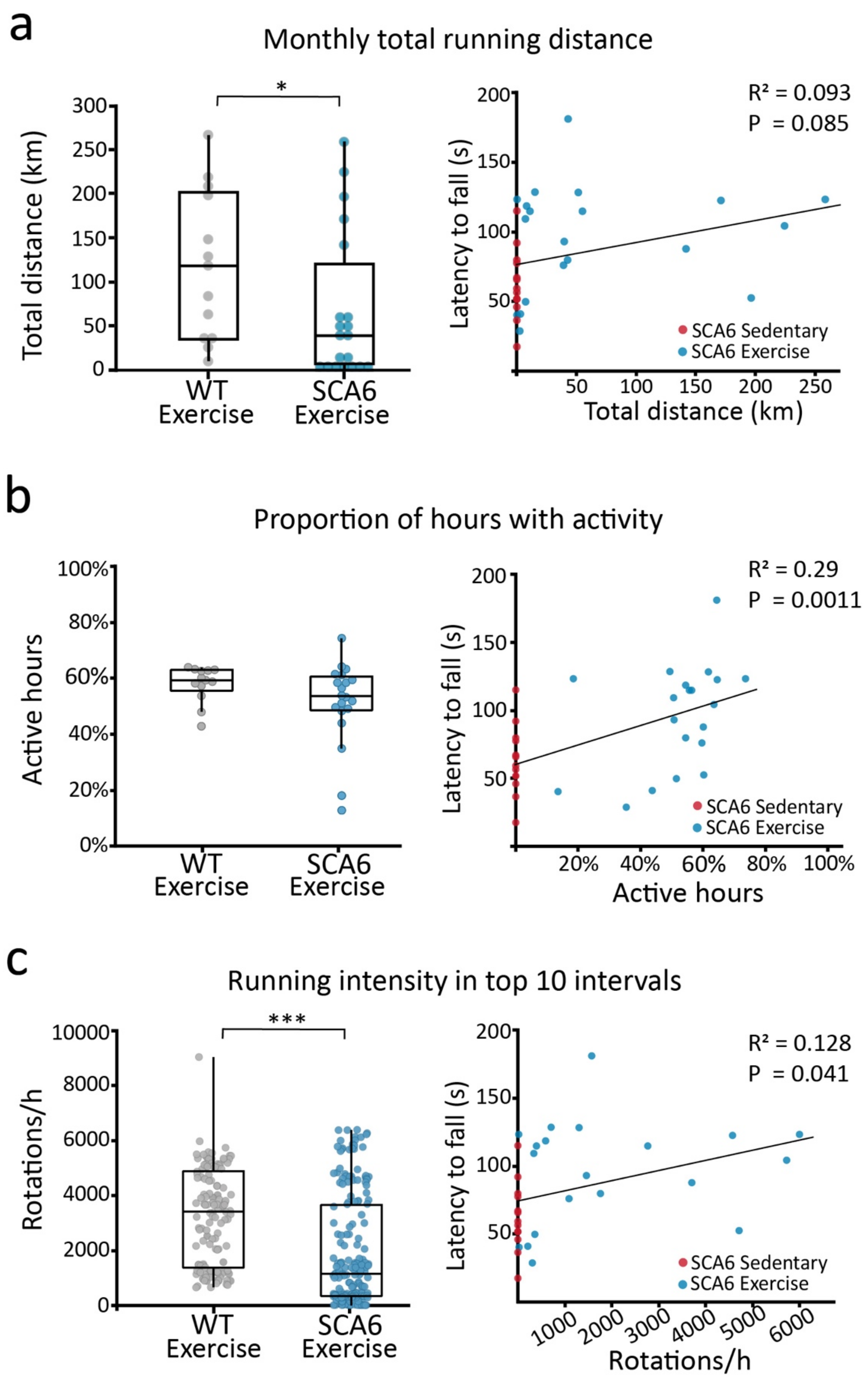


**Supplementary figure 2. SCA6$^{84Q/84Q}$ mice exercise with a lower intensity and over moderately shorter distances than WT mice.**

(**a**) Left: WT mice run slightly further than SCA6$^{84Q/84Q}$ mice over a month (significantly different, Wilcoxon rank test, P = 0.046). Right: There is no significant correlation observed between running distance and motor coordination, represented as latency to fall(s) on day 5 of rotarod testing. (**b**) Left: There is no difference between the proportion of hours/day that WT and SCA6$^{84Q/84Q}$ mice spent running (not significantly different, P = 0.093). Right: However, there is weak but significant correlation between the proportion of daily hours SCA6$^{84Q/84Q}$ mice spend running and their motor coordination. (**c**) Left: WT mice run with greater intensity than SCA6$^{84Q/84Q}$ mice, which is defined as the average speed reached in each mouses top 30-minute intervals that they spent running (P < 0.0001). Right: There is a weak but significant correlation between each mouse's top running intensity and motor coordination. Statistical comparisons made with Mann Whitney *U* test, N = 13 WT mice, N = 20 SCA6$^{84Q/84Q}$ mice.* P < 0.05, *** P < 0.001, not significant when not shown.

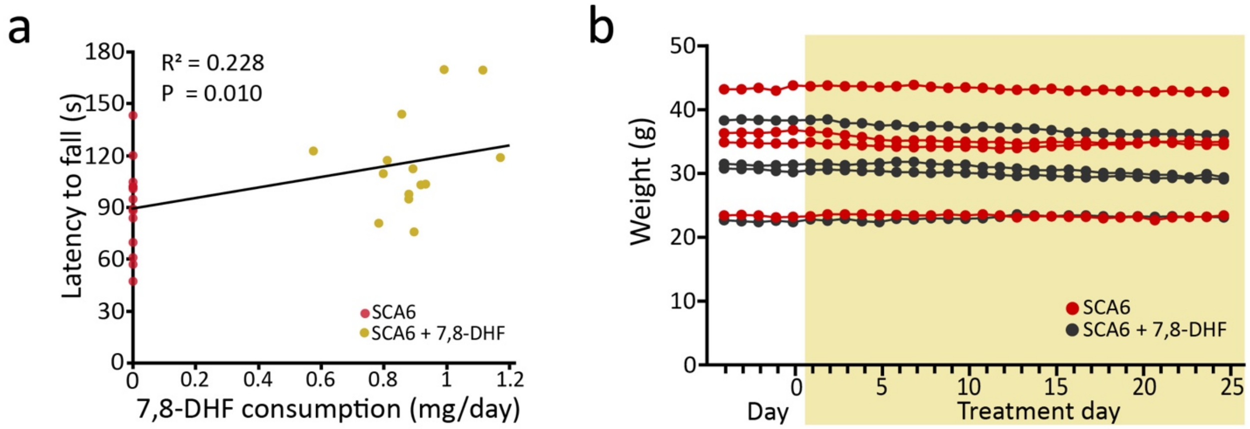


**Supplementary Fig. 3. 7,8-DHF treatment is well tolerated and consumption levels are positively correlated with rotarod performance in SCA6$^{84Q/84Q}$ mice.**

(**a**) We observe a weak correlation between average daily 7,8-DHF consumption and motor coordination (as measured by day 5 rotarod performance) in 7 month SCA6$^{84Q/84Q}$ mice treated for 1 month with 7,8-DHF or control (SCA6 N = 14, SCA6 + 7,8-DHF N = 14). (**b**) Mouse weight was not affected by 7,8-DHF consumption, suggesting that it is well tolerated. Individual data lines correspond to a subset of individual SCA6$^{84Q/84Q}$ mice that were either treated with 7,8-DHF (black) or control (red) weighed daily.

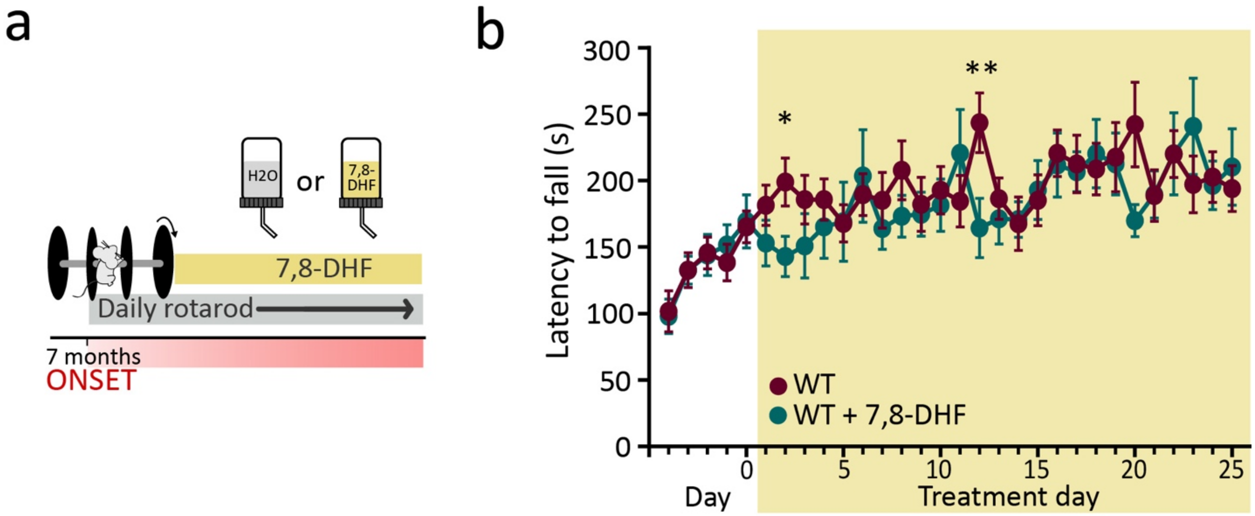


**Supplementary Fig. 4. Chronic administration of 7,8-DHF does not alter motor coordination in wildtype mice.**

(**a**) Schematic of experimental design. (**b**) WT mice treated with 7,8-DHF show no sustained, significant improvement compared to control WT mice. (WT N = 4, WT + DHF N = 4).

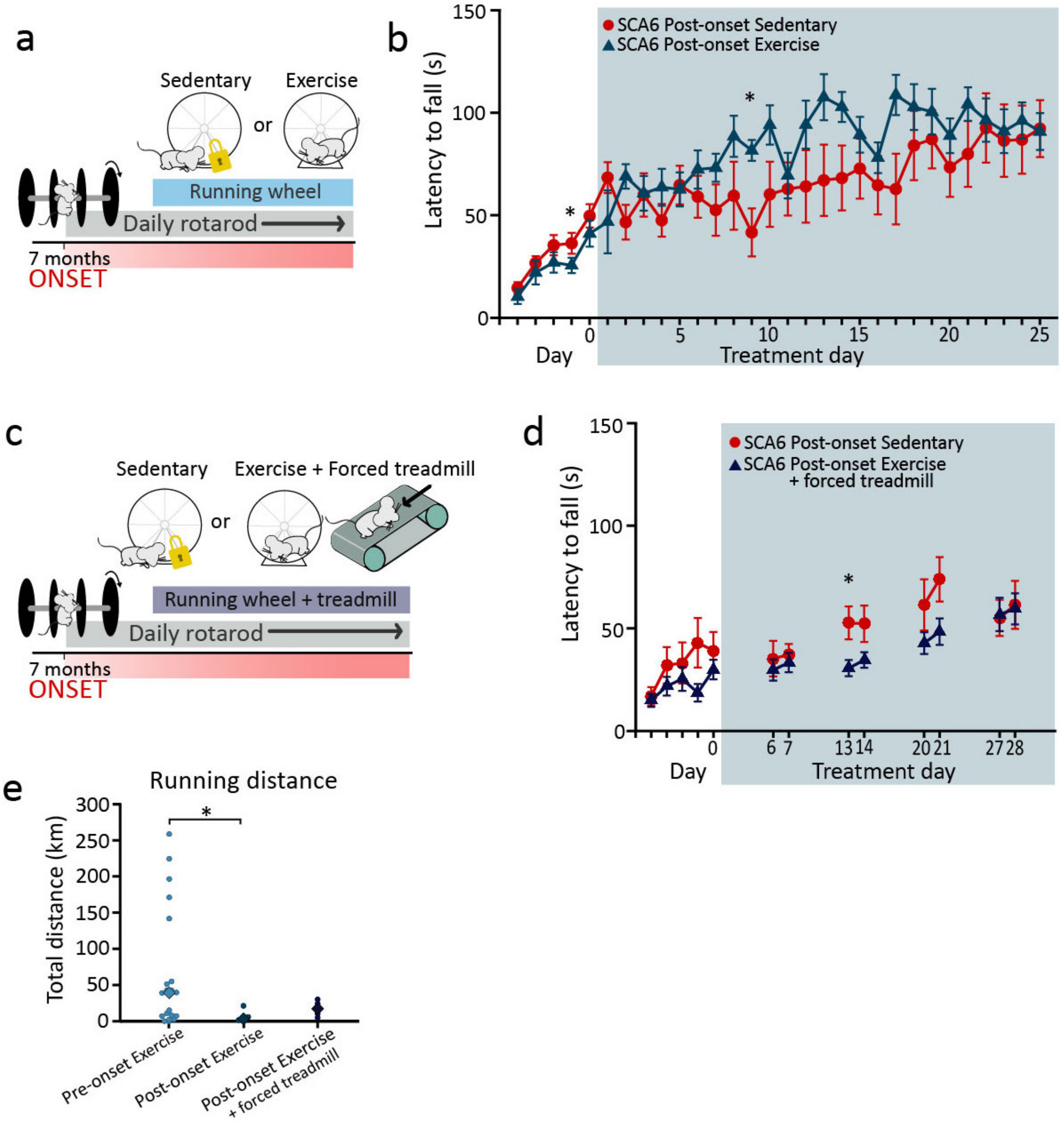


**Supplementary Fig. 5. Exercise treatment does not rescue motor deficits if it is started after the onset of ataxia**

(**a**) Schematic of exercise protocol and rotarod testing in mice after disease onset. (**b**) Post-onset SCA6$^{84Q/84Q}$ mice (SCA6) given running wheels (Exercise) performed no differently than Sedentary mice. (all days compared with Mann-Whitney *U* test, except days 2, 9, 16, 24 compared with Student's *t* test, all P values were >0.05 and so were not significantly different, except day 4, P = 0.028 and day 14, P = 0.028; SCA6$^{84Q/84Q}$ N = 5, SCA6$^{84Q/84Q}$ + Exercise N = 4). (**c**) To increase

Exercise by post-onset SCA6$^{84Q/84Q}$ mice, we used daily bouts of forced treadmill exercise in addition to voluntary wheel running. (**d**) However, we observed no significant improvement in motor coordination when mice underwent both forced and spontaneous Exercise (days 1, 2, 4, 5,18, 19, 25 were compared with Mann-Whitney *U* test, all other days compared with Student's *t* test, all P values were >0.05 and so were not significantly different, except day 18 p = 0.029; SCA6$^{84Q/84Q}$ N = 3, SCA6$^{84Q/84Q}$ forced exercise N = 4). (**e**) Total running distance over the 1 month period in SCA6$^{84Q/84Q}$ pre-onset (6 months), early onset (7 months) and early onset (7 months) with forced treadmill (One-way ANOVA followed by post-hoc Dunn's multiple comparison test, only SCA6$^{84Q/84Q}$ pre-onset and SCA6$^{84Q/84Q}$ early onset were significantly different, P = 0.038; SCA6$^{84Q/84Q}$ pre-onset vs SCA6$^{84Q/84Q}$ early onset with forced treadmill, P = 0.42, SCA6$^{84Q/84Q}$ early onset vs SCA6$^{84Q/84Q}$ early onset with forced treadmill, P = 0.11; SCA6$^{84Q/84Q}$ pre-onset N = 16, SCA6$^{84Q/84Q}$ early onset N = 5, SCA6$^{84Q/84Q}$ early onset with forced treadmill N = 4).

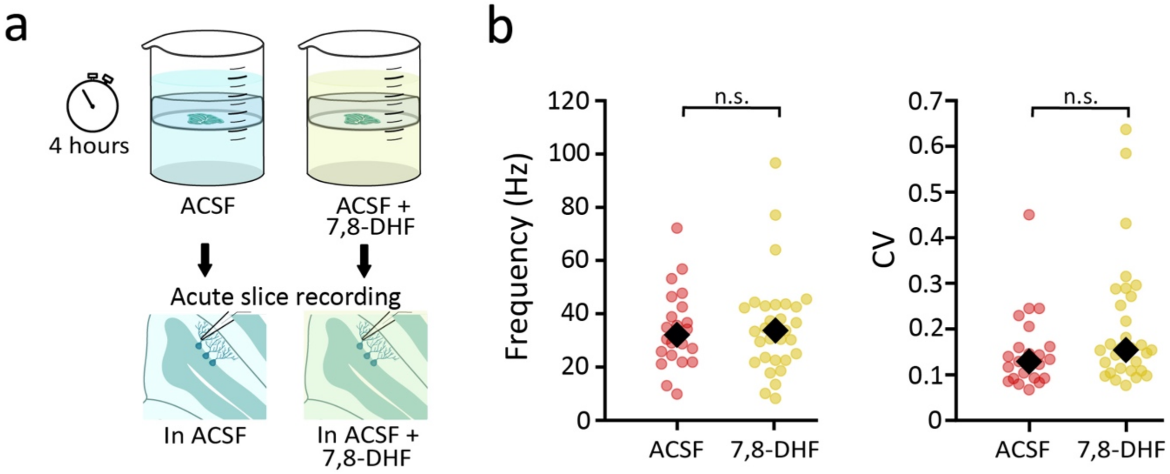


**Supplementary Fig. 6 Acute 7,8-DHF application did not alter Purkinje cell firing properties or running distance in SCA6$^{84Q/84Q}$ mice.**

(**a**) Schematic showing experimental design to test whether short-term (4 h) administration of 7,8-DHF was sufficient to alter Purkinje cell firing. (**b**) Neither Purkinje cell frequency (left) nor regularity (right, CV) was altered by 4 hours of 7,8-DHF treatment (not significantly different, Mann-Whitney *U* test, P = 0.91for frequency and P = 0.070 for CV; SCA6 control, N = 3 mice and n = 23 cells; SCA6 + acute DHF, N = 3 mice and n = 30 cells, n.s. P > 0.05).

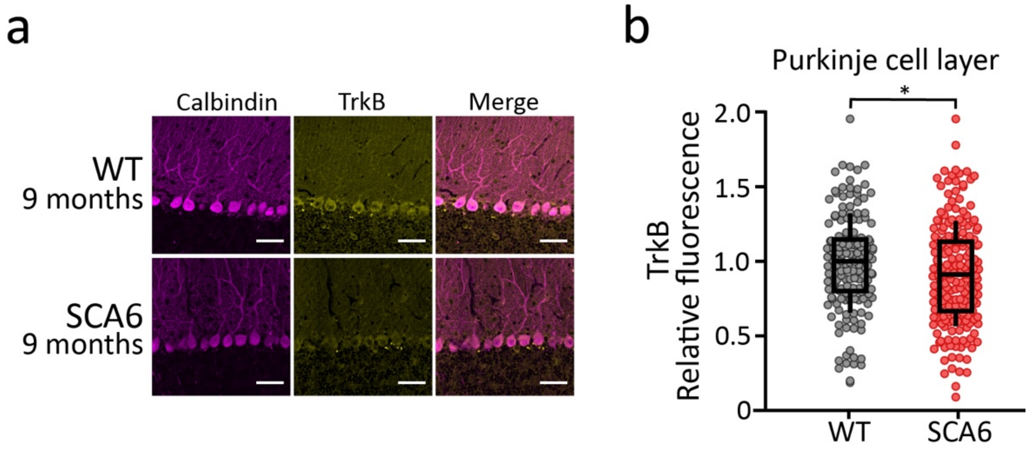


**Supplementary Fig. 7. TrkB receptors are still expressed in Purkinje cells of SCA6$^{84Q/84Q}$ mice at 9 months.**

(**a**) TrkB immunoreactivity is still observed in WT and SCA6$^{84Q/84Q}$ cerebellum from 9 month old mice (anti-calbindin is used to label Purkinje cells). Scale bar = 50 µm. (**b**) TrkB intensity is reduced in SCA6$^{84Q/84Q}$ mice at 9 months to a similar level as observed at 7 months (**Fig. 1**; significantly different, Mann-Whitney *U* test, P = 0.038; WT N = 3 mice, SCA6 N = 4 mice).

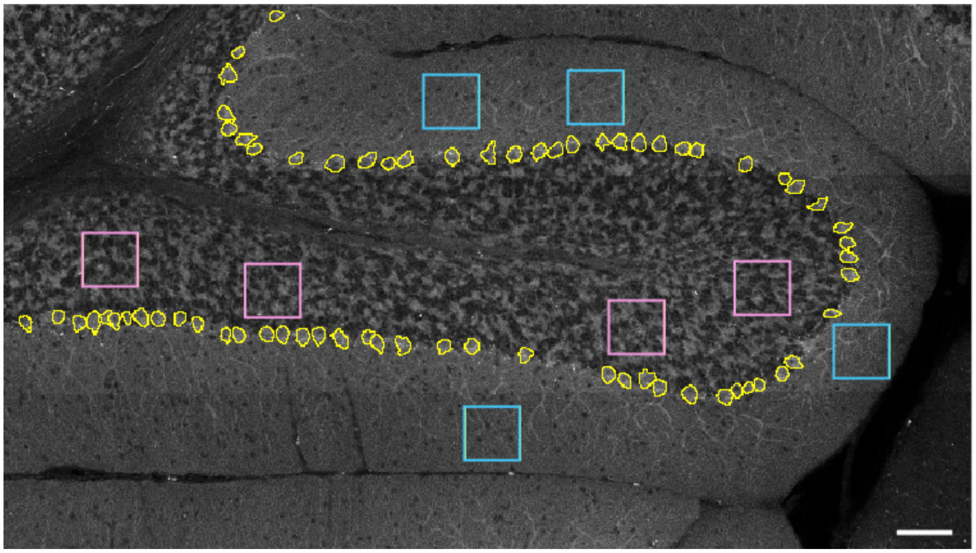

**Supplementary Fig. 8. Illustration of immunofluorescence analysis**

Purkinje cell somata (yellow outlines) are circled in the imaging channel expressing calbindin (typically the red channel). These circles are then used to measure the BDNF intensity in that channel (typically the green channel). Four regions of interest (ROIs) are randomly chosen in both the granule cell layer (pink boxes) and molecular cell layer (blue boxes) in the calbindin channel and measurements of BDNF intensity are taken in the BDNF channel. Scale bar, 75 μm.

| Antigen | Antibody | Supplier |
|---|---|---|
| Calbindin | Anti-calbindin (mouse) [300] | Swant |
| GFAP | Anti-GFAP (rat) [13-0300] | Thermo Fisher Scientific |
| BDNF | Anti-BDNF (rabbit) [ab108319] | ABCAM |
| TrkB | Anti-TrkB (rabbit) [ab9872] | Milipore |
| Secondary anti-mouse | Anti-mouse Alexa 594 [715-585-150] | Jackson immunoresearch |
| Secondary anti-rat | Anti-rat Alexa 594 [112-585-167] | Jackson immunoresearch |
| Secondary anti-rabbit | Anti-rabbit Alexa 488 [711-545-152] | Jackson immunoresearch |
| Secondary anti-mouse | Anti-mouse DyLight 405 [35500BID] | Thermo Fisher Scientific |

**Supplementary Table 1. List of antibodies used**
